## Supplementary Figure and Table for "Targeting resident astrocytes attenuates neuropathic pain after spinal cord injury"

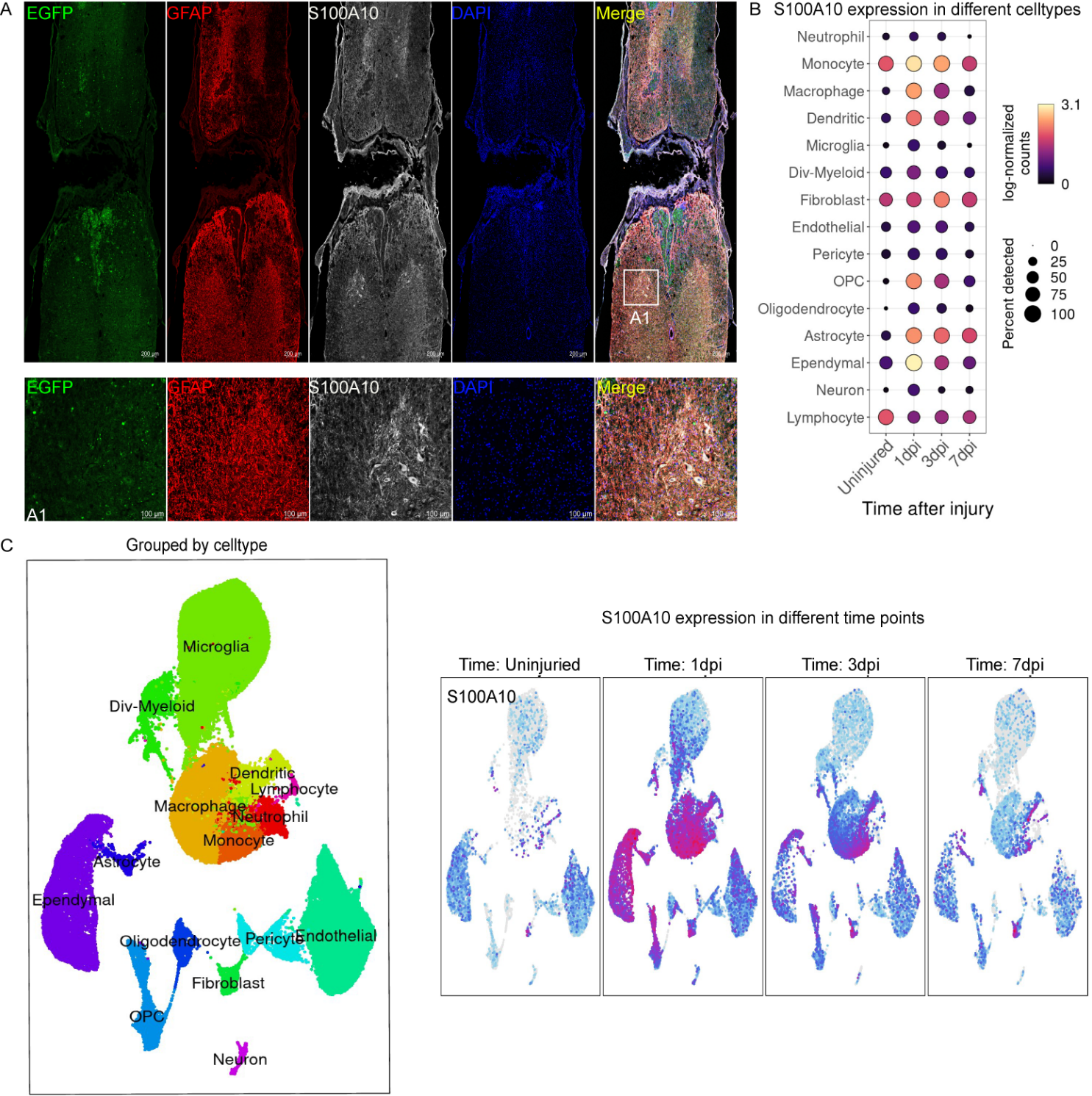

**Figure 1-figure supplement 1. S100 family protein p11 (S100A10)** **expressed by various cell types. (A)**, S100A10 expressed in astrocytes and neuronal like cells of gray matter. GFAP (red, marker of astrocytes), EGFP (green), S100A10 (white). n = 3 biological repeats. Scale bars had been indicated in pictures. (**B-C**), S100A10 expressed by various cell types at different time after SCI. Figures were analyzed and downloaded through online data from injured mouse spinal cords (https://jaeleelab.shinyapps.io/sci_singlecell/).

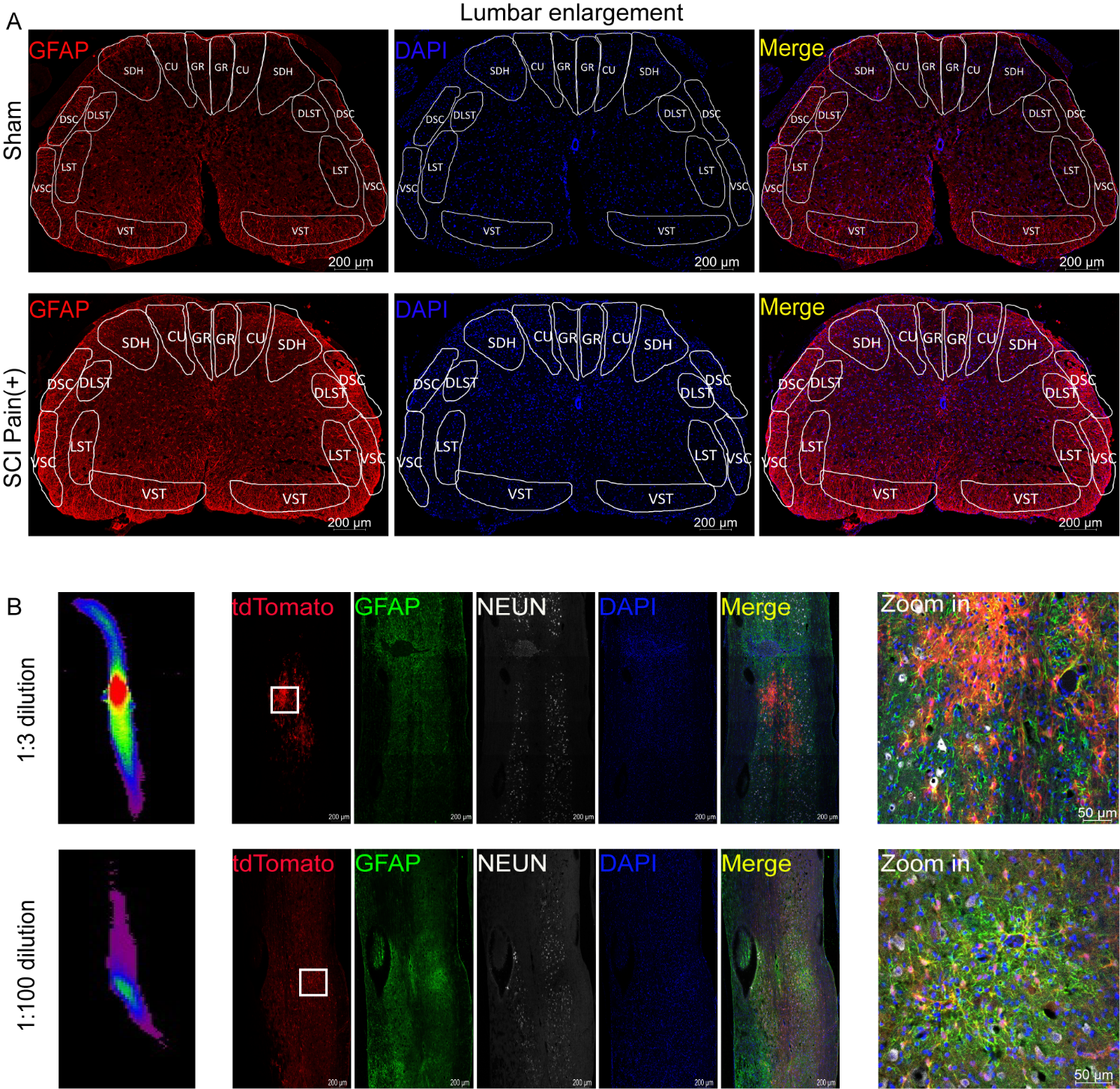
**Figure 2-figure supplement 1.** **Astrocytes activation in the lumbar enlargement of neuropathic pain mice. The results of AAV2/5-GfaABC1D-Cre targeted astrocytes in lumbar enlargement**. (**A**), images of immunofluorescent staining using GFAP (red) as marker of astrocytes. The location of three largest ascending tracts of lumbar enlargements, including (1) CU, GR; (2) LST, DLST, VST; (3) DSC, VSC, along with SDH and gray matter. CU=cuneate fasciculus, GR=gracile fasciculus, LST=lateral spinothalamic tract, DLST=dorsolateral spinothalamic tract, VST=ventral spinothalamic tract, DSC=dorsal spinocerebellar tract, VSC=ventral spinocerebellar tract, SDH=superficial dorsal horn. Scale bar, 200μm. (**B**), targeting astrocyte in lumbar enlargement with AAV2/5-GfaABC1D-Cre. 1:3 dilution, ≥ 0.33E+13 V.G/ml; 1:100 dilution, ≥1E+11 V.G/ml. Images of immunofluorescent staining using GFAP (green) and NEUN (white) as characteristic markers of neuron and astrocyte. n = 3 biological repeats. Scale bars had been indicated in pictures.

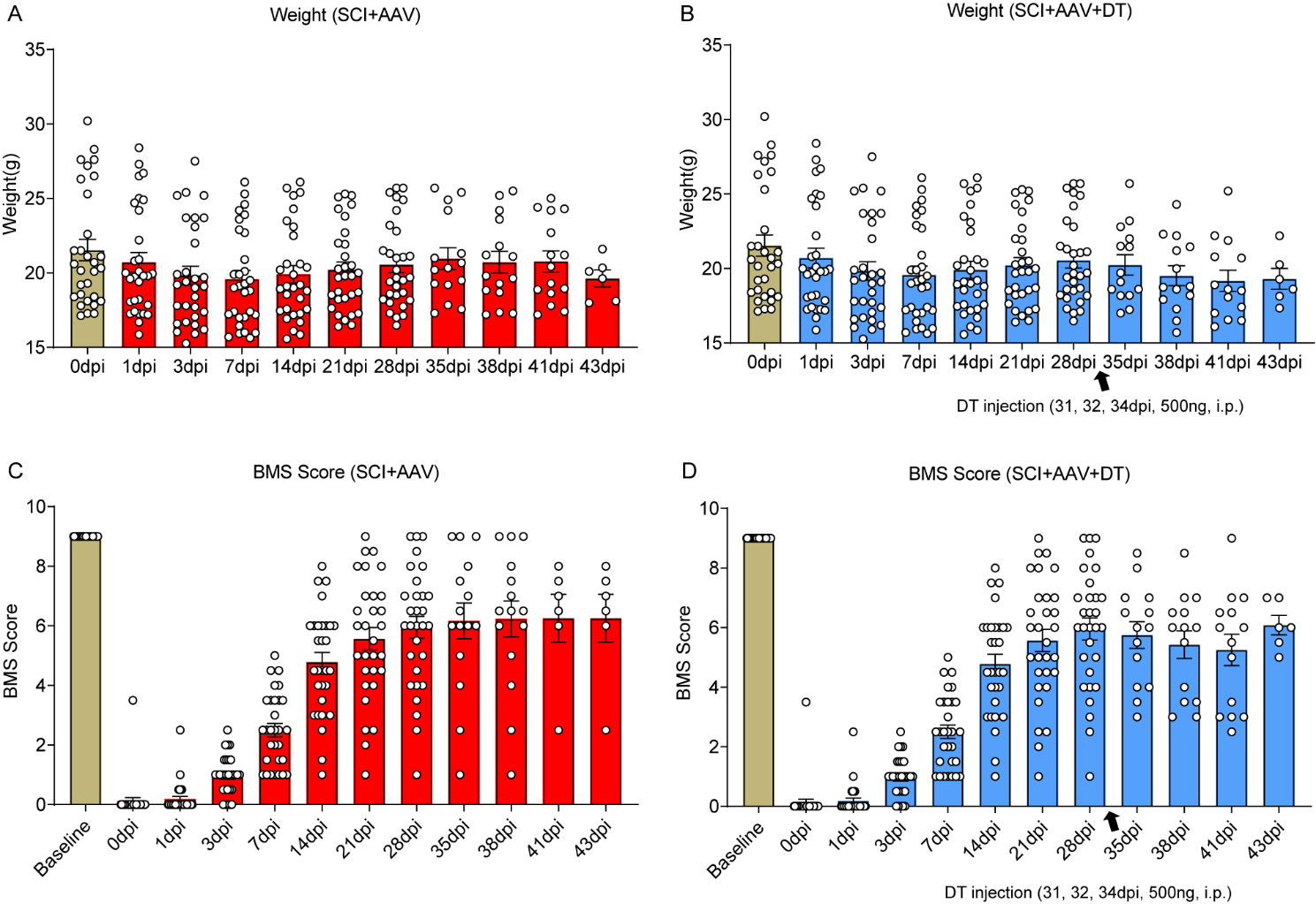
**Figure 3—figure supplement 1. Selective astrocyte elimination in lumbar enlargement attenuated neuropathic pain.**

Change of weight (**A-B**), BMS scores (**C-D**) in mice after astrocyte elimination. Sham group, n=30; sham+AAV group, n=30; SCI+AAV group, n=36; SCI+AAV+DT group, n=36. Arrow, 500 ng DT injection was performed on 31, 32 and 34 dpi (days post-injury).

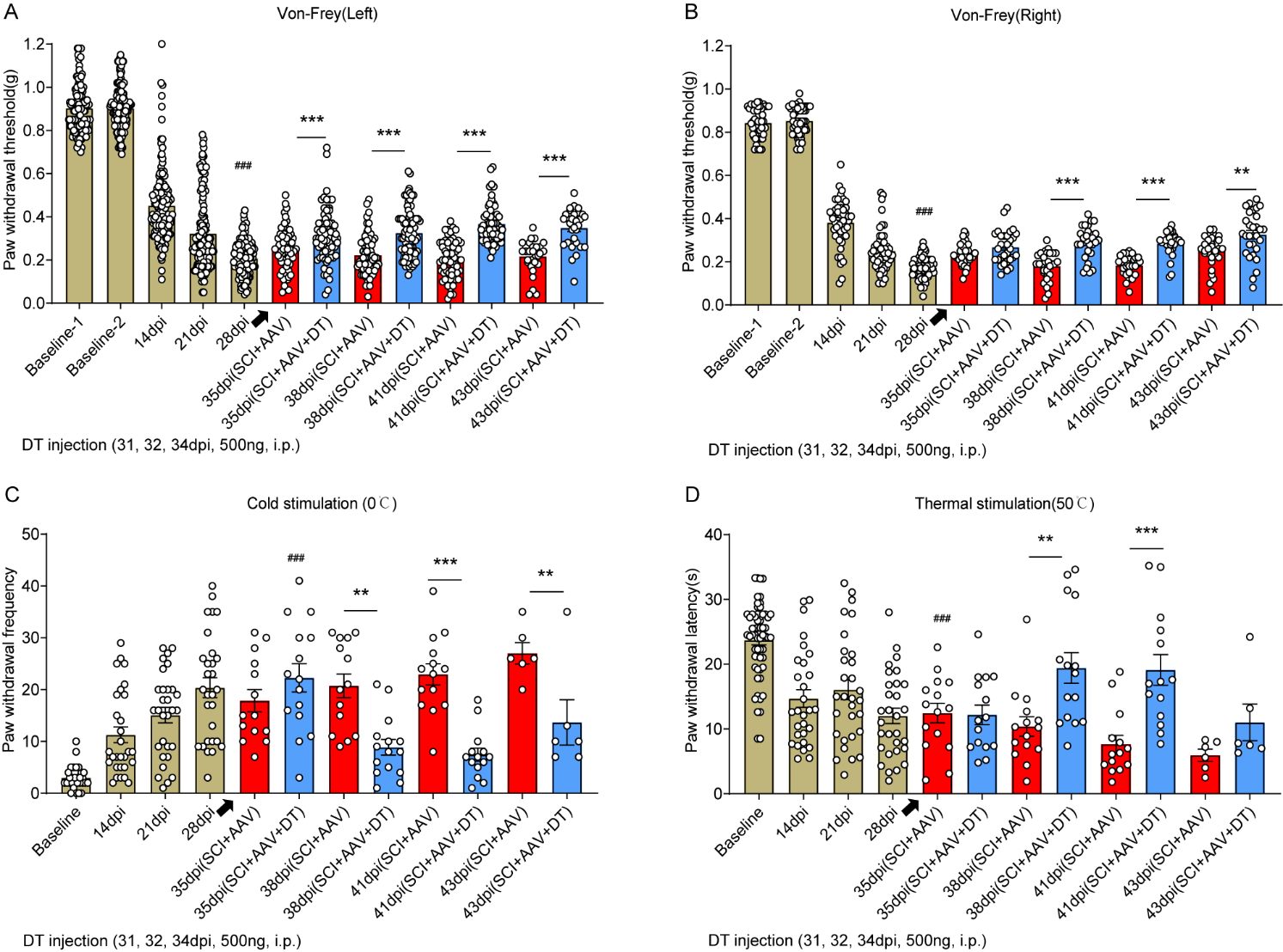
**Figure 3—figure supplement 2. Selective astrocyte elimination in lumbar enlargement attenuated neuropathic pain.**

Change of mechanical allodynia (**A-B**), cold hyperalgesia (**C**) and thermal hyperalgesia (**D**) in mice after astrocyte elimination. SCI+AAV group, n=36; SCI+AAV+DT group, n=36. Arrow, 500 ng DT injection was performed on 31, 32 and 34 dpi (days post-injury). Left, left hindlimbs; right, right hindlimbs. i.p., intraperitoneal injection. Statistical significance was determined by two-way ANOVA followed by Student Newman–Keuls post hoc test. **P < 0.01, ***P < 0.001, SCI+AAV+DT group vs. SCI+AAV group.

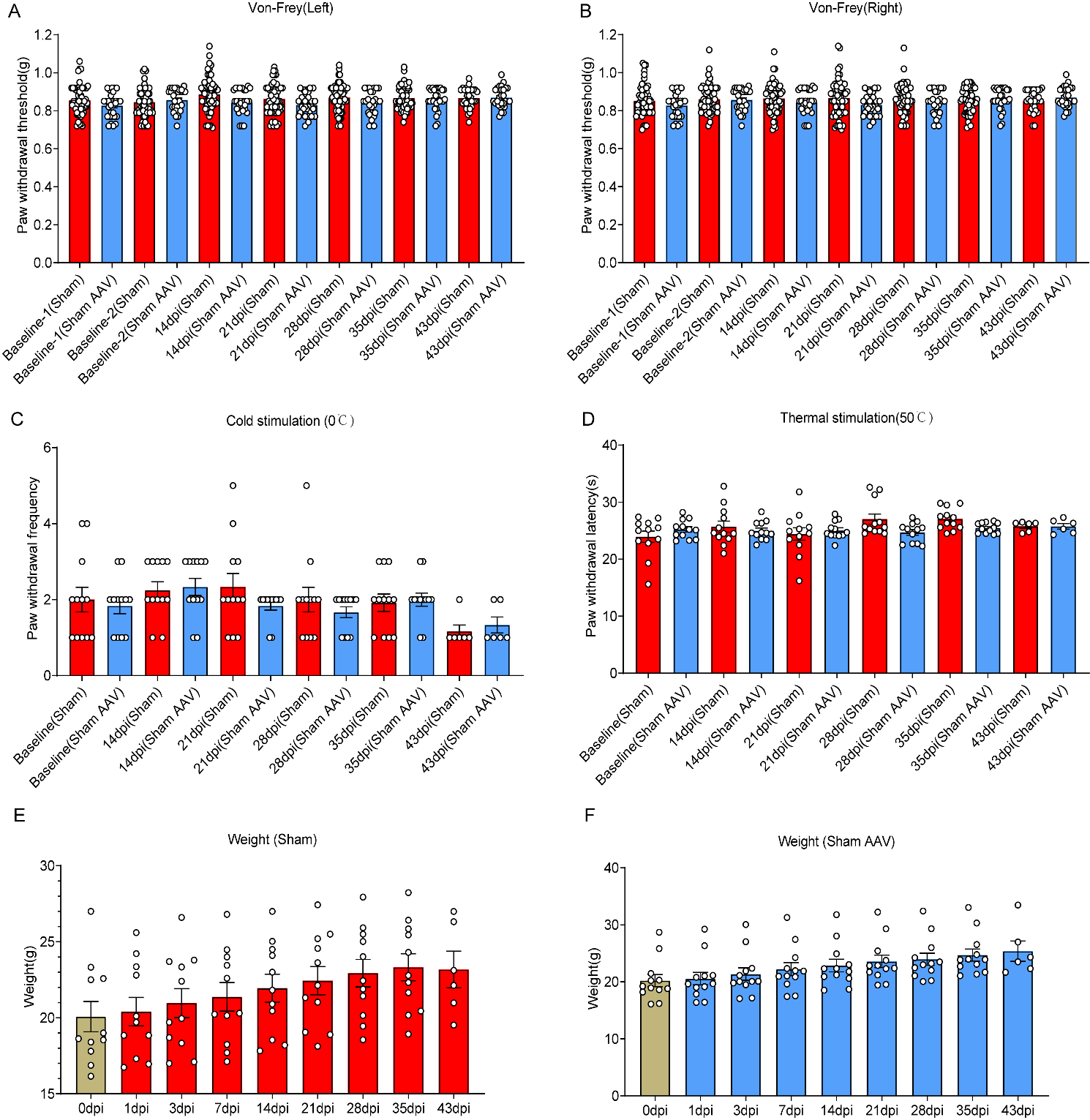
**Figure 3—figure supplement 3. Selective astrocyte elimination in lumbar enlargement attenuated neuropathic pain.**

Change of mechanical allodynia (**A-B**), cold hyperalgesia (**C**) and thermal hyperalgesia (**D**), weight (**E-F**) in mice of sham group and sham+AAV group. Left, left hindlimbs; right, right hindlimbs.

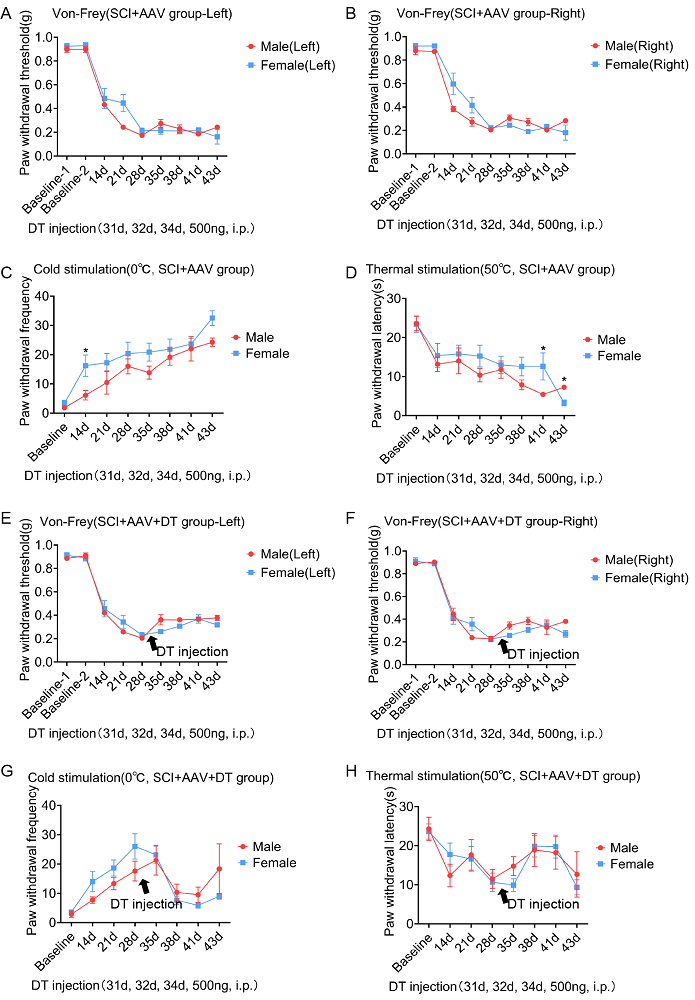

**Figure 3-figure supplement 4.** **No differences in neuropathic pain were observed between male and female mice.** (A-H), Change of mechanical allodynia (A-B, E-F), cold hyperalgesia (C, G) and thermal hyperalgesia (D, H) in mice of SCI+AAV group (n=36) and SCI+AAV+DT group (n=36). Arrow, 500 ng DT injection was performed on 31, 32 and 34 dpi (days post-injury). Left, left hindlimbs; right, right hindlimbs. Values are the mean ± SEM. Statistical significance was determined by one-way ANOVA followed by Student Newman–Keuls post hoc test. *P < 0.05.

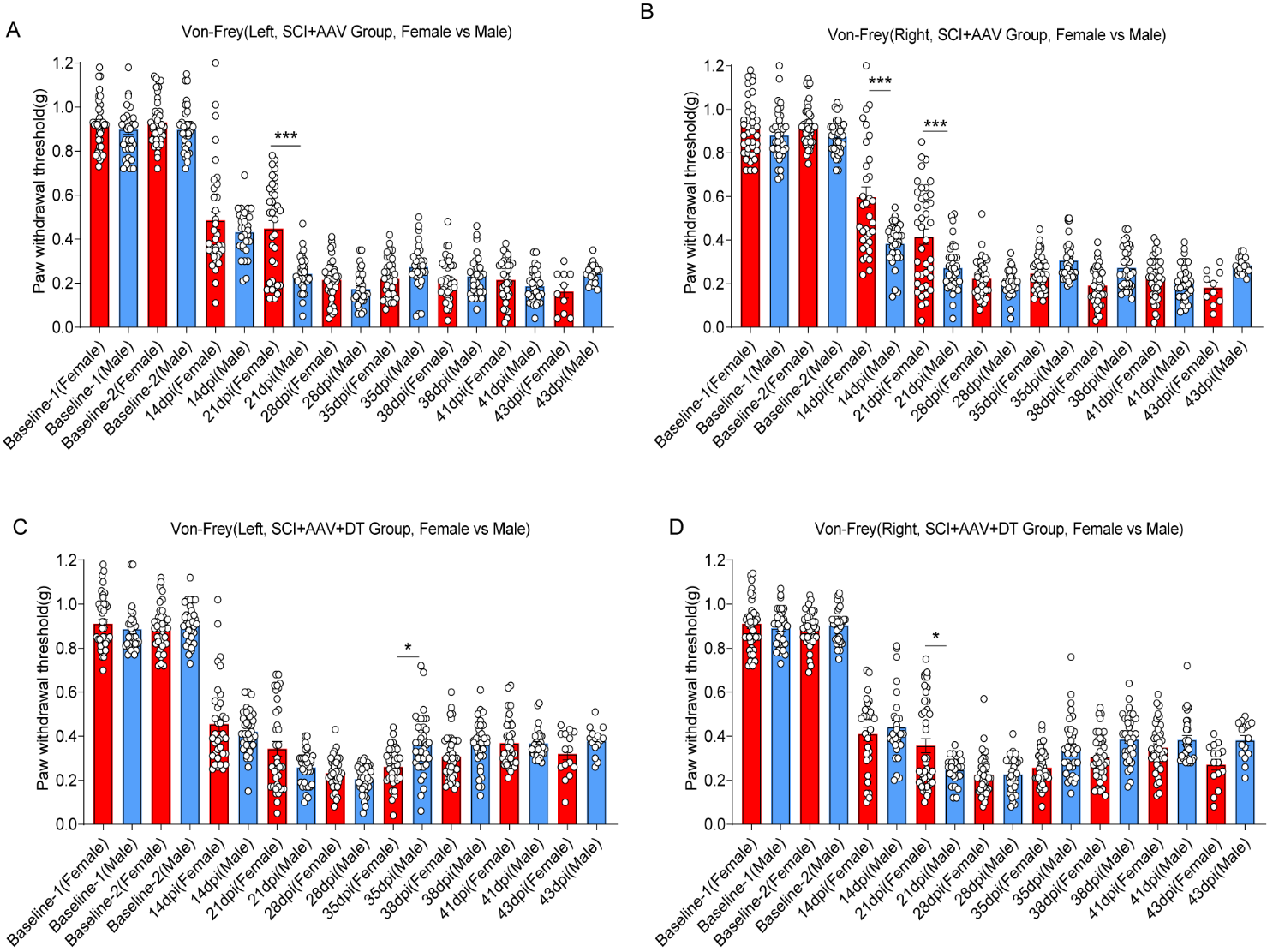
**Figure 3—figure supplement 5. No significant differences in neuropathic pain were observed between male and female mice.**

Change of mechanical allodynia (**A-D**) in mice of SCI+AAV group and SCI+AAV+DT group after astrocyte elimination. SCI+AAV group, n=36; SCI+AAV+DT group, n=36. Left, left hindlimbs; right, right hindlimbs. Statistical significance was determined by one-way ANOVA followed by Student Newman–Keuls post hoc test. *P < 0.05, ***P < 0.001, Female mice vs. male mice.

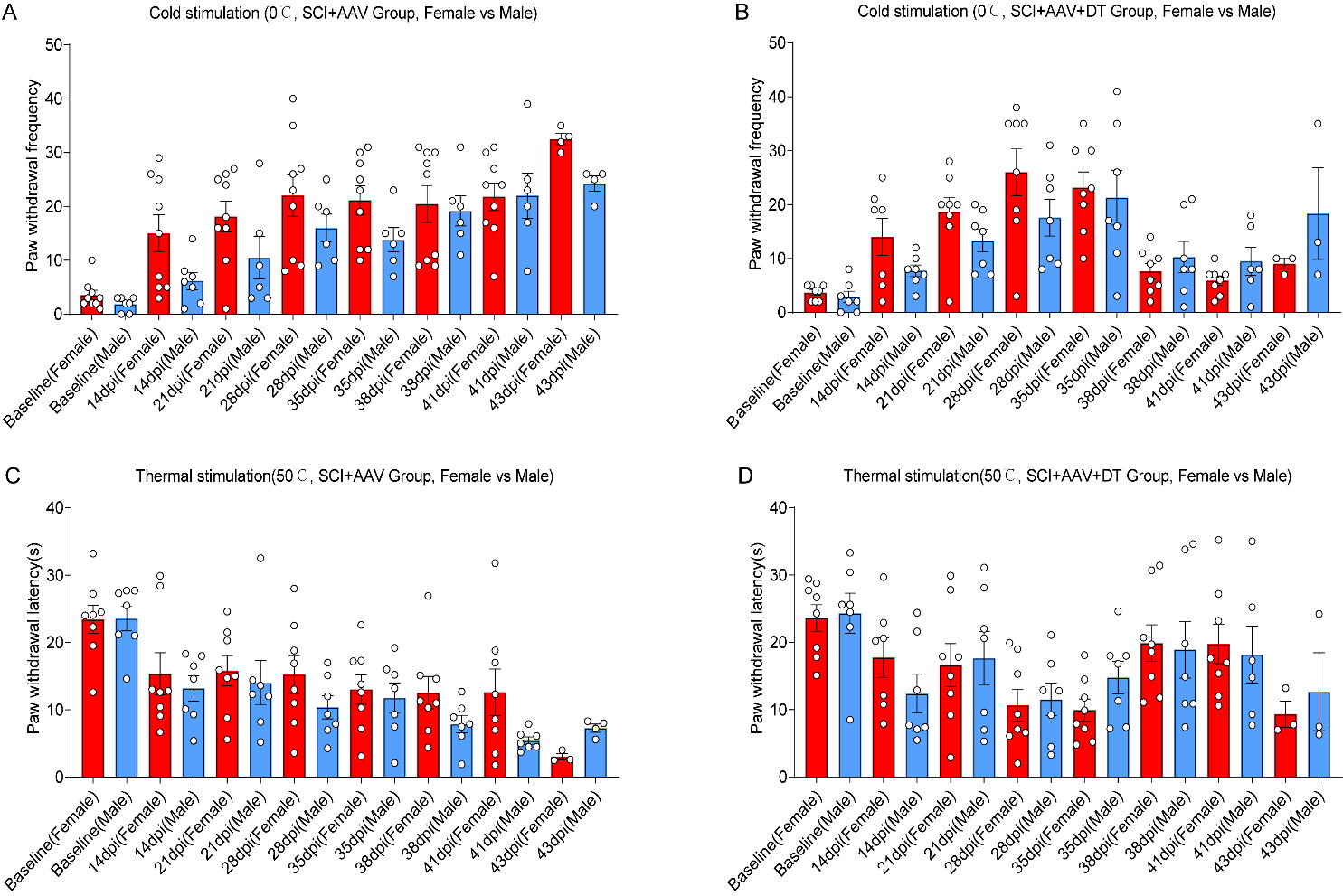
**Figure 3—figure supplement 6. No significant differences in neuropathic pain were observed between male and female mice.**

Change of cold hyperalgesia (**A-B**) and thermal hyperalgesia (**C-D**) in mice of SCI+AAV group and SCI+AAV+DT group after astrocyte elimination.

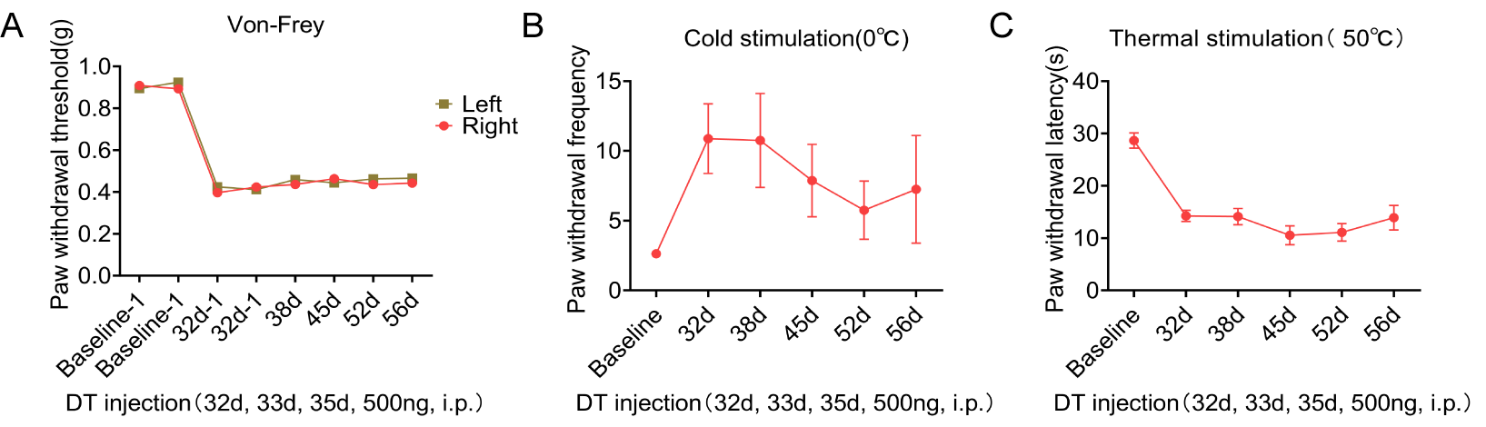

**Figure 3-figure supplement 7.** **Pain symptoms were similar in wild-type mice with or without DT injection.** Mechanical allodynia (**A**), cold (**B**) and thermal (**C**) hypersensitivity test results after DT injection in wild-type mice. n = 8 biological repeats. Values are the mean ± SEM. Statistical significance was determined by two-way ANOVA was performed followed by Student Newman–Keuls post hoc test.

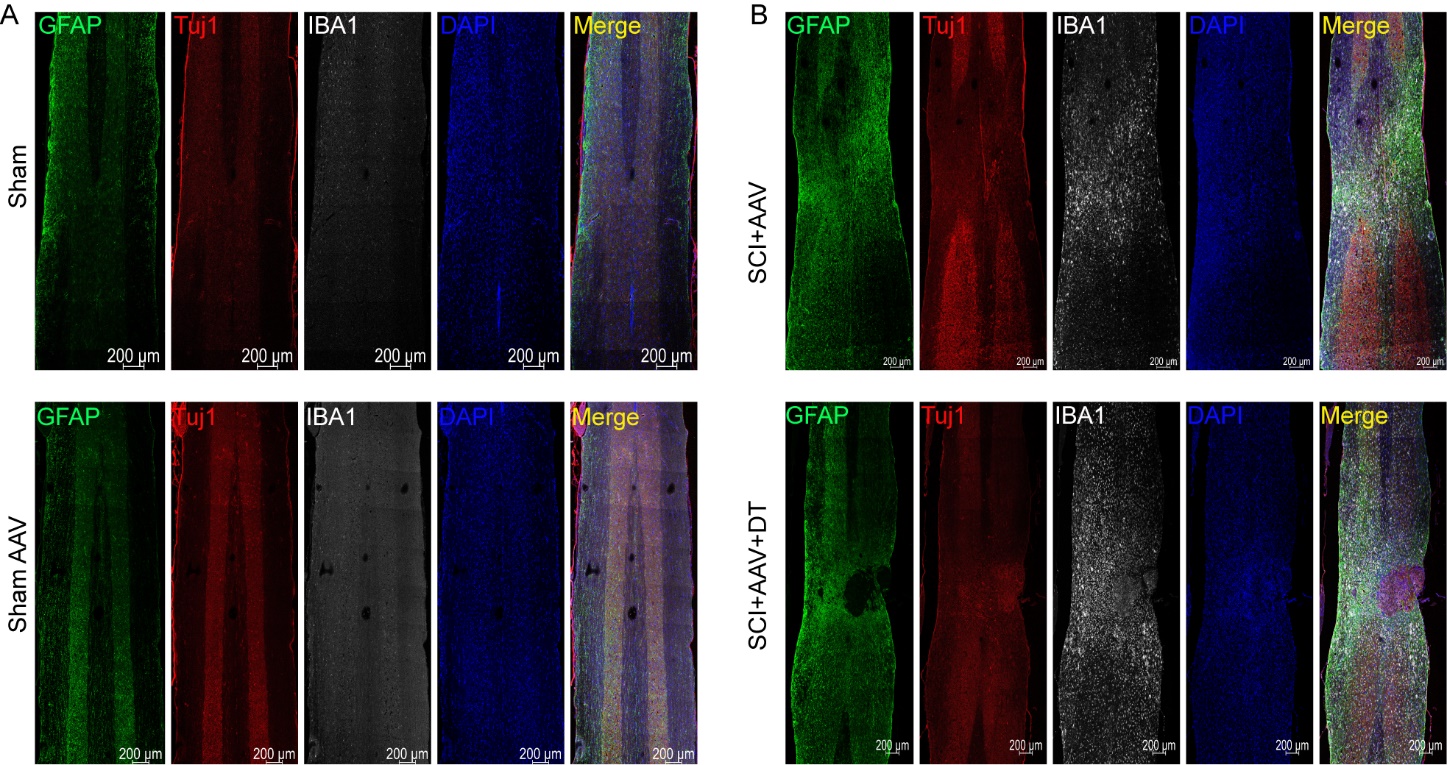
**Figure 4-figure supplement 1.** **Full fluorescence images of lesion area.** (**A-B**), the full fluorescence images of lesion area. Images of immunofluorescent staining using GFAP (green), Tuj1 (red) and IBA1 (white) as characteristic markers of astrocyte, neuron and microglia, respectively. n = 3 biological repeats. Scale bar, 200μm.

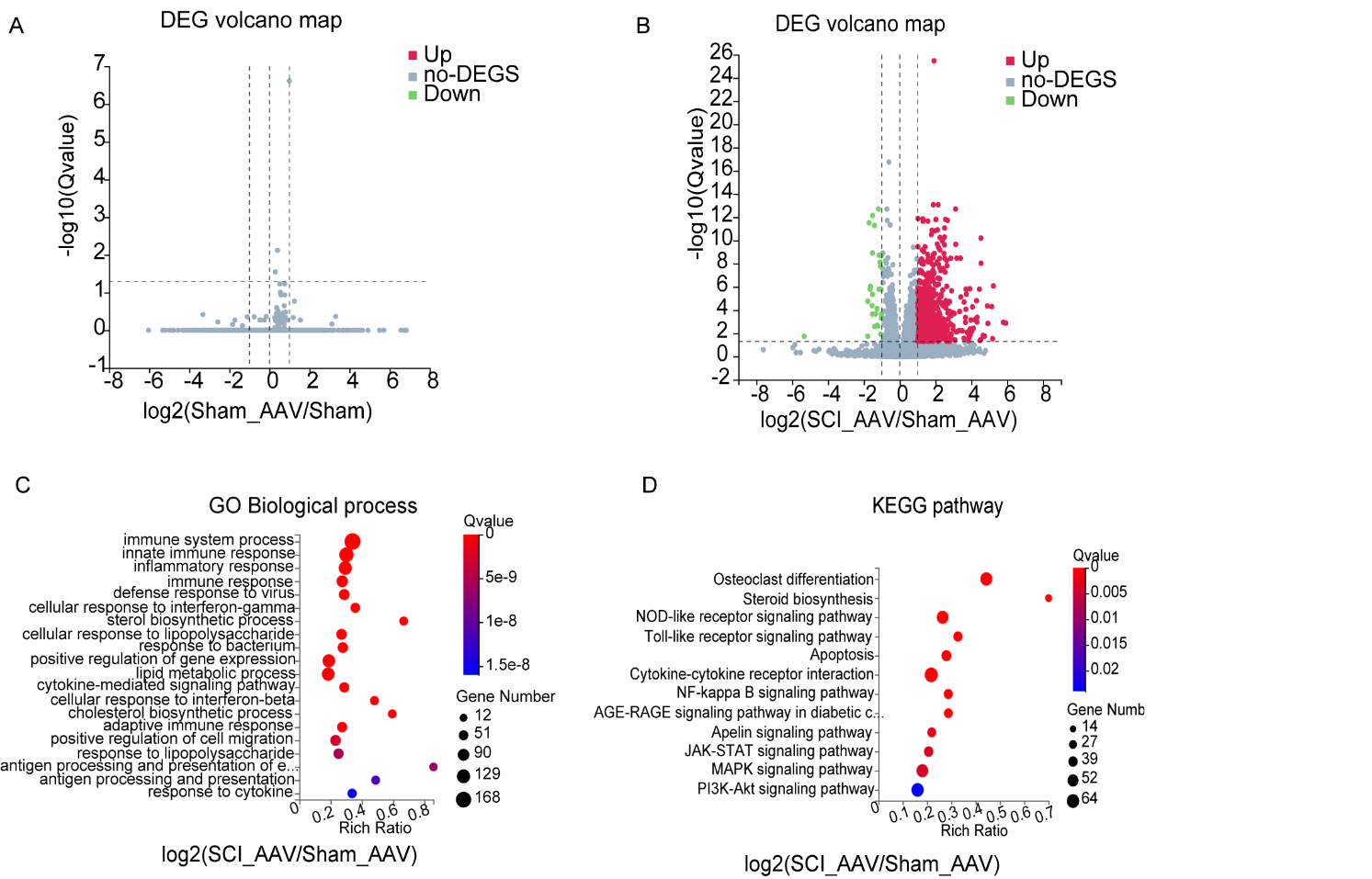
**Figure 5-figure supplement 1.** **RNA-seq analysis results between SCI+AAV group and sham AAV group.** (**A-B**), Volcano maps showed the DEGs between groups. (**C-D**), the GO enrichment and KEGG pathway analysis between the SCI+AAV group and sham AAV group. n = 3 biological repeats.

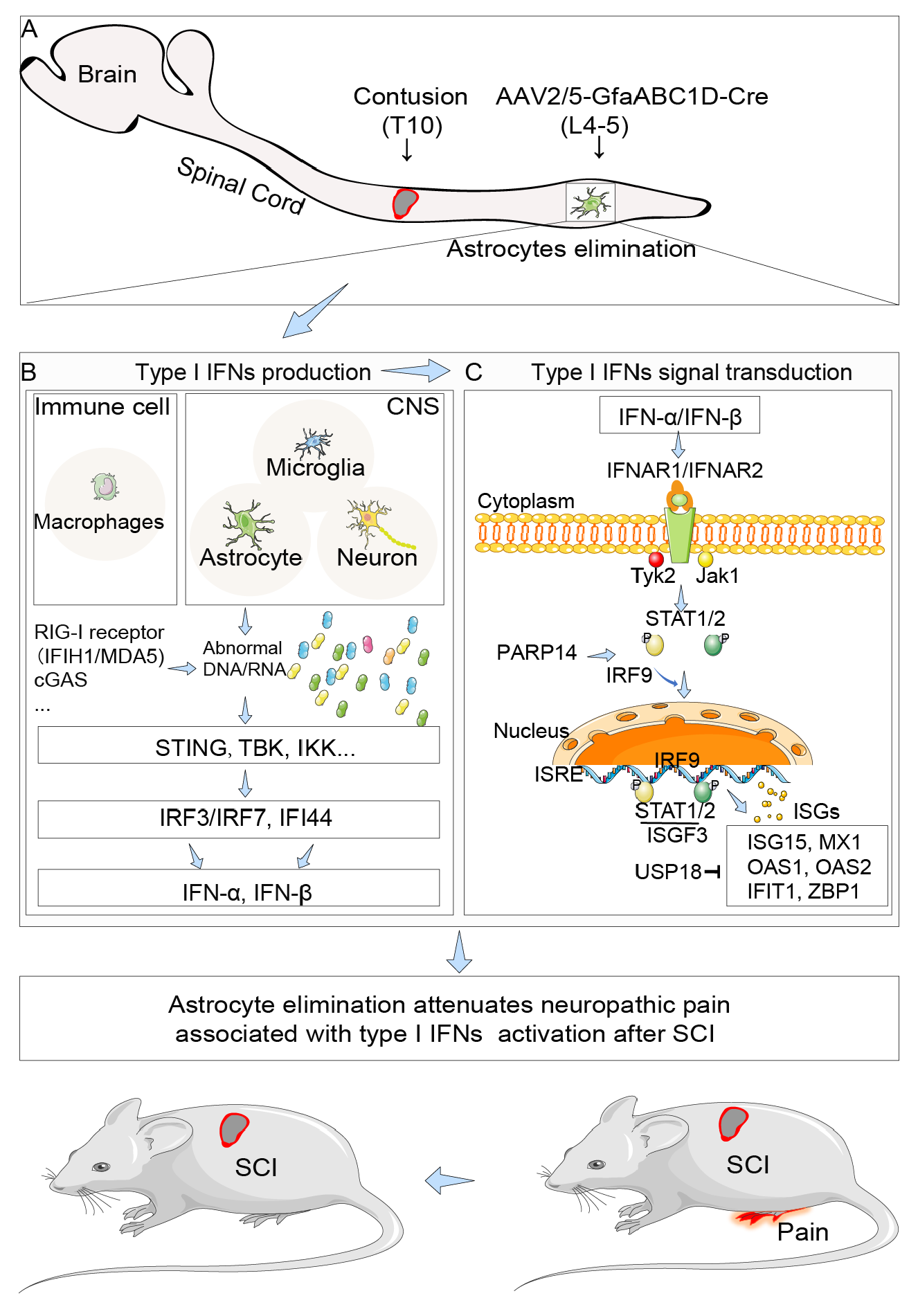
**Figure 7—figure supplement 1. The potential schematic diagram of selective resident astrocytes elimination attenuated neuropathic pain after SCI**. (**A**), astrocytes in lumbar enlargement were targeted and selectively eliminated through transgenic mice injected with an adeno-associated virus vector (AAV2/5-GfaABC1D-Cre) and diphtheria toxin. Selective astrocyte elimination in lumbar enlargement could attenuate neuropathic pain after SCI, which were associated with type I IFNs signal and microglia activation. (**B**), the production of type I IFNs. Type I IFNs production is mainly caused by the contact of innate immune cells (mainly macrophages, microglia and astrocyte in CNS) surface or internal receptors (RIG-I receptor Ifih1/Mda5, Cgas, etc.) with virus specific antigenic substances (DNA, RNA), then through intracellular signal molecule transmission (*Sting, Tbk, Ikk*, etc.), and finally activate the transcription factor IRF3/7 to promote the expression of type I IFNs, including IFN-α and IFN-β. (**C**), the signal transduction of type I IFNs. Type I IFNs bind to the same two membrane spanning polypeptide chains type I IFNs receptor 1/2 (IFNAR1/2) (Borden *et al.*, 2007; Owens *et al.*, 2014) and lead to cross phosphorylation and activation of Tyk2/JaK1. Activation of Tyk2 and JAK1 phosphorylate STAT1/2 to form a heterodimer, which then translocate to the nucleus and associate with IRF9 to further form the heterotrimeric transcription factor complex IFN-stimulated gene factor-3 (ISGF3). Finally, ISGF3 translocate to the nucleus and bind to specific IFN-response elements (ISREs) to control the expression of IFNs-stimulated genes (ISGs) (Rothhammer *et al.*, 2016), including *Isg15, Mx1, Oas1, Oas2, Ifit1, Zbp1*.

Supplementary File 1a. Transgenic mice and genotyping primers

| Mouse | Cat# | Resource | Genotyping primers (5'to3') |
| --- | --- | --- | --- |
| C57BL/6-Foxj1^em1(GFP-CreERT2-polyA)Smoc^**^§^** | NM-KI-200133 | Model Organisms Centre, Shanghai | F1 \| GTTTGGGCCTTCCTACCCTC; R1 \| TTCGAGATGTGCACGACGAT; F2 \| TCTTTCCTCTCGGGGTAGGG; R2 \| CTTGTAGTTGCCGTCGTCCT |
| FVB-Tg(GFAP-cre)^25Mes/J #^ | JAX:004600 | The Jackson Laboratory (JAX) | F \| ACT CCT TCA TAA AGC CCT; R \| ATC ACT CGT TGC ATC GAC CG |
| Tg(CAG-Dre) *^Smoc*^* | NM-TG-00026 | Model Organisms Centre, Shanghai | F \| ACTCCTTGCCGATGTTCCTCAG; R \| TTGTCCCAAATCTGGCGGAG |
| Gt(ROSA)26Sor^em1(CAG-LSL-RSR-tdTomato-2A-DTR)Smoc &^ | NM-KI-190086 | Model Organisms Centre, Shanghai | F1 \| TCAGATTCTTTTATAGGGGACACA; R1 \| TAAAGGCCACTCAATGCTCACTAA; F2 \| ATGAAGCTGCTGCCGTCGG; R2 \| TCAGTGGGAATTAGTCATGCCCAA |
| Ai9(LSL-tdTomato) - B6.Cg-Gt(ROSA) 26Sor ^tm9(CAG-tdTomato) Hze^/^J^ **^†^** | JAX:007909 | The Jackson Laboratory (JAX) | F1 \| AAGGGAGCTGCAGTG GAGTA; R1 \| CCGAAAATCTGTGGGAAGTC; F2 \| GGCATTAAAGCAGCGTATCC­; R2 \| CTGTTCCTGTACGGCATGG |
| **^§^** C57BL/6-Foxj1^em1(GFP-CreERT2-polyA)Smoc^ mice are under the control of promoters with specific expression in ependymal cells (FoxJ1), which have been widely used for identification of cells that express Foxj1 in the central nervous system and peripheral organs. These mice are on R26R Cre-reporter background and tamoxifen administration will lead to permanent and heritable expression of the reporter gene EGFP.  ^#^ FVB-Tg(GFAP-cre)^25Mes/J^ is a tool mouse expressed Cre recombinase under the control of the human glial fibrillary acidic protein promoter (GFAP). Cre recombinase recognizes loxP site and can mediate recombination. | | | |
| ^*^ Tg(CAG-Dre) *^Smoc^* is a tool mouse expressed Dre recombinase driven by extensive CAG promoter. Dre Rox is similar to Cre loxP system. Dre recombinase recognizes Rox site and can mediate chromosome recombination. | | | |
| ^&^ Gt(ROSA)26Sor^em1(CAG-LSL-RSR-tdTomato-2A-DTR)Smoc^ is a reporter mouse expressed loxP site (loxP-Stop- loxP), rox site (Rox-Stop-Rox), reporter gene tdtomato and diphtheria toxin receptor (DTR) driven by extensive CAG promoter. | | | |
| **^†^** Ai9(LSL-tdTomato) - B6.Cg-Gt(ROSA) 26Sor ^tm9(CAG-tdTomato) Hze^/^J^ is a reporter mouse expressed tdtomato. | | | |

Supplementary File 1b. Primers for type I interferon signal pathway genes

| Primer name | Primers (5'to3') | |
| --- | --- | --- |
| Ifit1 | forward | *5’-CTGAGATGTCACTTCACATGGAA-3’* |
|  | reverse | *5’-GTGCATCCCCAATGGGTTCT-3’* |
| Irf7 | forward | *5’-GAGACTGGCTATTGGGGGAG-3’* |
|  | reverse | *5’-GACCGAAATGCTTCCAGGG-3’* |
| Irf3 | forward | *5’-GAGAGCCGAACGAGGTTCAG-3’* |
|  | reverse | *5’-CTTCCAGGTTGACACGTCCG-3’* |
| Ifi44 | forward | *5’-AACTGACTGCTCGCAATAATGT-3’* |
|  | reverse | *5’-GTAACACAGCAATGCCTCTTGT-3’* |
| Isg15 | forward | *5’-GGTGTCCGTGACTAACTCCAT-3’* |
|  | reverse | *5’-TGGAAAGGGTAAGACCGTCCT-3’* |
| Usp18 | forward | *5’-TTGGGCTCCTGAGGAAACC-3’* |
|  | reverse | *5’-CGATGTTGTGTAAACCAACCAGA-3’* |
| Oas2 | forward | *5’-TTGAAGAGGAATACATGCGGAAG-3’* |
|  | reverse | *5’-GGGTCTGCATTACTGGCACTT-3’* |
| Oas3 | forward | *5’-TCTGGGGTCGCTAAACATCAC-3’* |
|  | reverse | *5’-GATGACGAGTTCGACATCGGT-3’* |
| Zbp1 | forward | *5’-AAGAGTCCCCTGCGATTATTTG-3’* |
|  | reverse | *5’-TCTGGATGGCGTTTGAATTGG-3’* |
| Dhx58 | forward | *5’-GGAAGTGATCTTACCTGCTCTGG-3’* |
|  | reverse | *5’-TTGCCTCTGTCTACCGTCTCT-3’* |
| Ifih1 | forward | *5’-AGATCAACACCTGTGGTAACACC-3’* |
|  | reverse | *5’-CTCTAGGGCCTCCACGAACA-3’* |
| Stat2 | forward | *5’-TCCTGCCAATGGACGTTCG-3’* |
|  | reverse | *5’-GTCCCACTGGTTCAGTTGGT-3’* |
| Irf9 | forward | *5’-GCCGAGTGGTGGGTAAGAC-3’* |
|  | reverse | *5’-GCAAAGGCGCTGAACAAAGAG-3’* |
| Parp14 | forward | *5’-TGCCAAGCAGTCAGTGATGTC-3’* |
|  | reverse | *5’-CCTGGAAAACTGTGTGCTCTAT-3’* |
| Eif2ak2 | forward | *5’-ATGCACGGAGTAGCCATTACG-3’* |
|  | reverse | *5’-TGACAATCCACCTTGTTTTCGT-3’* |
| Stat1 | forward | *5’-TCACAGTGGTTCGAGCTTCAG-3’* |
|  | reverse | *5’-GCAAACGAGACATCATAGGCA-3’* |
| GAPDH | forward | *5’-AGTGCCAGCCTCGTCTCATA-3’* |
|  | reverse | *5’-TGAACTTGCCGTGGGTAGAG-3’* |

| Supplementary File 1c. Primers for pro- and anti-inflammatory microglial marker genes | | |
| --- | --- | --- |
| Primer name | Primers (5'to3') | |
| TNF-α | forward | 5′-ATGCTGGGACAGTGACCTGG-3′ |
|  | reverse | 5′-CCTTGATGGTGGTGCATGAG-3′ |
| IL-1β | forward | 5′-CCAAAAGATGAAGGGCTGCT-3′ |
|  | reverse | 5′-TCATCAGGACAGCCCAGGTC-3′ |
| IL-6 | forward | 5′-GAGAGCCGAACGAGGTTCAG-3′ |
|  | reverse | 5′-GAAGGCCGTGGTTGTCACC-3′ |
| iNOS | forward | 5′-GTTCTCAGCCCAACAATACAAGA-3′ |
|  | reverse | 5′-GTGGACGGGTCGATGTCAC-3′ |
| IL-4 | forward | 5′-CACGGATGCGACAAAAATCA-3′ |
|  | reverse | 5′-CTCGTTCAAAATGCCGATGA-3′ |
| TGFβ1 | forward | 5′-TGGAGCTGGTGAAACGGAAG-3′ |
|  | reverse | 5′-ACAGGATCTGGCCACGGAT-3′ |
| IL-10 | forward | 5′-GCTCTTACTGACTGGCATGAG-3′ |
|  | reverse | 5′-CGCAGCTCTAGGAGCATGTG-3′ |
| Arg1 | forward | 5′-CTCCAAGCCAAAGTCCTTAGAG-3′ |
|  | reverse | 5′-AGGAGCTGTCATTAGGGACATC-3′ |

| Supplementary File 1d. Primers for Type I IFN signal related genes | | |
| --- | --- | --- |
| Primer name | Primers (5'to3') | |
| *IRF7* | forward | 5′-*GAGACTGGCTATTGGGGGAG*-3′ |
|  | reverse | 5′-*GACCGAAATGCTTCCAGGG*-3′ |
| *IRF3* | forward | 5′-*GAGAGCCGAACGAGGTTCAG*-3′ |
|  | reverse | 5′-*CTTCCAGGTTGACACGTCCG*-3′ |
| *ISG15* | forward | 5′-*GGTGTCCGTGACTAACTCCAT*-3′ |
|  | reverse | 5′-*TGGAAAGGGTAAGACCGTCCT*-3′ |
| *STAT2* | forward | 5′-*TCCTGCCAATGGACGTTCG*-3′ |
|  | reverse | 5′-*GTCCCACTGGTTCAGTTGGT*-3′ |
| *IRF9* | forward | 5′-*GCCGAGTGGTGGGTAAGAC*-3′ |
|  | reverse | 5′-*GCAAAGGCGCTGAACAAAGAG*-3′ |
| *STAT1* | forward | 5′-*TCACAGTGGTTCGAGCTTCAG*-3′ |
|  | reverse | 5′-*GCAAACGAGACATCATAGGCA*-3′ |
| *IFNb* | forward | 5′-*CAGCTCCAAGAAAGGACGAAC*-3′ |
|  | reverse | 5′-*GGCAGTGTAACTCTTCTGCAT*-3′ |
